# Whole genome structural predictions reveal hidden diversity in putative oxidative enzymes of the lignocellulose degrading ascomycete *Parascedosporium putredinis* NO1

**DOI:** 10.1101/2023.08.08.552407

**Authors:** Conor JR Scott, Daniel R Leadbeater, Nicola C Oates, Sally R James, Katherine Newling, Yi Li, Nicholas GS McGregor, Susannah Bird, Neil C Bruce

## Abstract

Economic valorisation of lignocellulose is paramount to realising a true circular bioeconomy; however, this requires the development of systems and processes to expand the repertoire of bioproducts beyond current renewable fuels, chemicals, and sustainable materials. *Parascedosporium putredinis* NO1 is an ascomycete that thrived at the later stages of a wheat-straw composting community culture, indicating a propensity to degrade recalcitrant lignin-enriched biomass, but exists within an underrepresented and underexplored fungal lineage. This strain has proven an exciting candidate for the identification of new enzymes targeting recalcitrant components of lignocellulose following the recent discovery of a new lignin β-ether linkage cleaving enzyme.

The first genome for the genus *Parascedosporium* for *P. putredinis* NO1 genome was sequenced, assembled, and annotated. The genome is 39 Mb in size, consisting of 21 contigs annotated to contain 9.998 protein-coding sequences. The carbohydrate-active enzyme (CAZyme) repertoire was compared to 2570 ascomycete genomes and in detail with *Trichoderma reesei*, *Fusarium oxysporum,* and sister taxa *Scedosporium boydii.* Significant expansion in the oxidative auxiliary activity class of CAZymes was observed in the *P. putredinis* NO1 genome resulting from increased sequences encoding putative lytic polysaccharide monooxygenases (LPMOs), oxidative enzymes acting within LPMO redox systems, and lignin-degrading laccases. *P. putredinis* NO1 scored above the 95^th^ percentile for AA gene density across the ascomycete phylum, suggesting a primarily oxidative strategy for lignocellulose breakdown. Novel structure-based searching approaches were employed, revealing 17 new sequences with structural similarity to LPMO, laccase, and peroxidase sequences and which are potentially new lignocellulose-degrading enzymes.

**Importance:** An annotated reference genome has revealed *P. putredinis* NO1 as a useful resource for the identification of new lignocellulose degrading enzymes for biorefining of woody plant biomass. Utilising a ‘structure-omics’ based searching strategy, new potentially lignocellulose-active sequences were identified that would have been missed by traditional sequence searching methods. These new identifications, alongside the discovery of novel enzymatic functions from this underexplored lineage with the recent discovery of a new phenol oxidase that cleaves the main structural β-O-4 linkage in lignin from *P. putredinis* NO1 highlights the underexplored and poorly represented family Microascaceae as particularly interesting candidates worthy of further exploration toward the valorisation of high value biorenewable products.

## Background

Energy consumption continues to grow rapidly alongside improvements in living standards, and fossil fuels continue to play a major role in industrial and agricultural sectors. With their widely accepted environmentally damaging effects, the need to move away from the use of fossil fuels and towards a net zero carbon fuel source is ever more pressing. Lignocellulosic residues consisting of cellulose, hemicellulose and lignin with minor amounts of pectins and nitrogen compounds offer the largest source of biomass for liquid fuel, chemicals and energy (1). However, biorefining of lignocellulose has so far been limited by the recalcitrant nature of the intricate and insoluble lignin network (2, 3).

Fungi are exceptional wood-degraders and are predominantly used to produce an array of bioproducts, including commercial enzyme cocktails used in biological processing of lignocellulosic biomass. Ascomycetes, known as soft-rot fungi, degrade lignocellulose by penetration of plant secondary cell walls with hyphae that secrete complex enzyme cocktails in abundance at the site of attack (4). *Parascedosporium putredinis* NO1 is a soft-rot ascomycete identified previously as dominant in the later stages of a mixed microbial compost community grown on wheat straw (5). This behaviour suggests that the fungus can efficiently deconstruct and potentially metabolise the more recalcitrant carbon sources in the substrate. Indeed, the recent discovery of a new oxidase enzyme that cleaves the major β-ether units in lignin in the *P. putredinis* NO1 secretome, which releases the pharmaceutically valuable compound tricin from wheat straw while simultaneously enhancing digestibility of the biomass (5), promotes a requirement for further exploration of this taxa.

Here, an annotated reference genome for *P. putredinis* NO1 reveals a repertoire of carbohydrate-active enzymes (CAZymes) and oxidative enzymes focused on degrading the most recalcitrant components of lignocellulose. Comparisons across the ascomycete tree of life suggest an increased proportion of oxidative enzymes within the CAZyme repertoire of *P. putredinis* NO1. Further investigation through CAZyme repertoire comparison with two other industrially relevant wood-degrading ascomycetes; *Trichoderma reesei*, and *Fusarium oxysporum*, as well as sister taxa *Scedosporium boydii* reveals expansion in families of enzymes with roles in the oxidative dissolution of lignocellulose and demonstrated this fungus to be an exciting candidate for the identification of new lignocellulose degrading activities.

Novel approaches were used to search the *P. putredinis* NO1 genome for potentially unannotated enzyme sequences with relation to three types of classic oxidative lignocellulose degraders: lytic polysaccharide monooxygenases (LPMOs), laccases, and peroxidases. Predicted structures were obtained for >96 % of the protein coding sequences in the genome. Structural searches were found to be effective at identifying multiple sequences for potentially novel proteins involved in lignocellulose breakdown which had low levels of structural similarity to the classic oxidative lignin and crystalline cellulose degrading enzymes. These sequences were also missed by sequence and domain-oriented searches. Further investigation and comparison of structures revealed varying levels of structural overlap despite the lack of sequence similarity. This strategy of combining search approaches can be adopted to identify divergent enzyme sequences which may have alternate lignocellulose degrading activity, variation in substrate-specificity, and different temperature and pH optima.

Further investigation and characterisation of such lignocellulose-degrading enzymes adds to the wealth of enzymes which can be incorporated into commercial enzyme cocktails to improve their effectiveness and boost the efficiency at which biomass is converted to renewable liquid fuel and value-added chemicals.

## Results and Discussion

### The Genome of *P. putredinis* NO1 Suggests a Strategy to Degrade the Most Recalcitrant Components of Lignocellulose

The *P. putredinis* NO1 genome was sequenced using nanopore sequencing with the Oxford Nanopore Technologies’ (ONT) MinION system to avoid errors in the assembly and annotation of coding regions resulting from long regions of repetitive DNA in eukaryotic genomes (6). The genome is 39 Mb in size and the assembly consists of 21 contigs, containing 9998 protein coding sequences. To investigate the lignocellulose-degrading enzyme repertoire of the *P. putredinis* NO1 genome, all protein coding sequences were annotated for CAZyme domains using the dbCAN server (7). In total, 795 CAZyme domains were predicted in the *P. putredinis* NO1 genome and the distribution of these domains across the CAZyme classes can be seen in **Figure 1A**. Glycoside Hydrolases (GH) are the most abundant CAZyme class with 290 identified. While Auxiliary Activities (AA) also make a large contribution with 162 domains. Glycosyl Transferases (GT) contribute 113 domains, and Carbohydrate Esterases (CE) contribute 51. Polysaccharide Lyases (PL) contribute the fewest with only 18 domains. In addition to these catalytic classes, 161 Carbohydrate Binding Modules (CBMs) were also identified.

**Figure 1.**
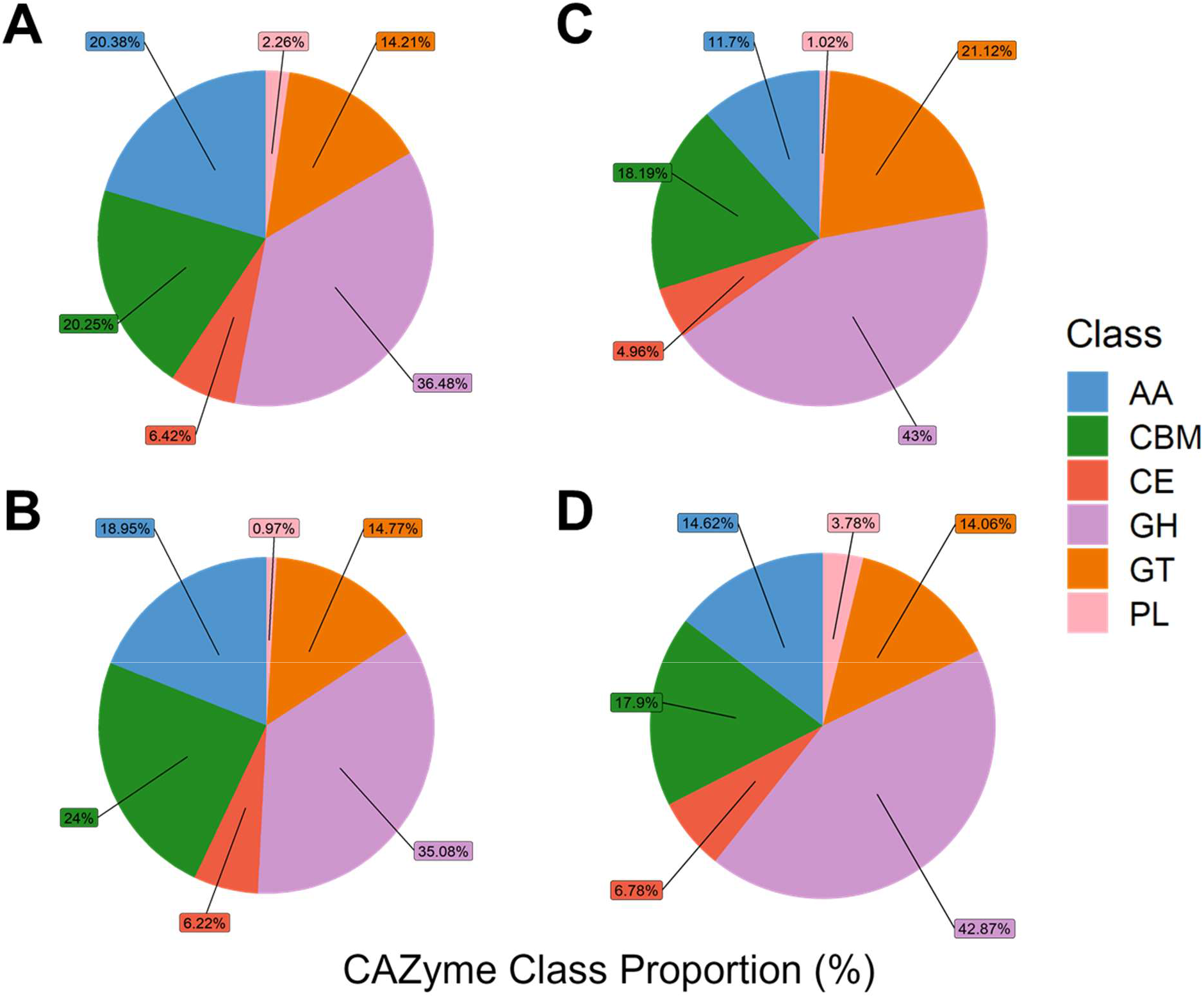
Comparison of CAZyme class repertoire. The number of CAZyme domains of each class for four lignocellulose degrading ascomycetes (**A**). The proportions of each class of CAZyme contributing to CAZyme repertoire for *P. putredinis* NO1 (**B**), *S. boydii* (**C**), *T. reesei* (**D**), and *F. oxysporum* (**E**). Auxiliary Activity (AA), Carbohydrate Binding Module (CBM), Carbohydrate Esterase (CE), Glycoside Hydrolase (GH), Glycosyl Transferase (GT), Polysaccharide Lyase (PL).

An interesting observation was the seemingly high number of AA class CAZymes observed in the *P. putredinis* NO1 genome. To investigate this further and more broadly within the scope of the ascomycete tree of life, CAZyme profiles of all available ascomycete genomes were elucidated (**Figure 2**). It was clear that *P. putredinis* NO1 has one of the highest proportions of AA class CAZymes within its repertoire among ascomycete fungi. *P. putredinis* NO1 (*Hypocreomycetidae*) belonged above the 95^th^ percentile for AA gene density among the highest AA populated genomes (25.59%), behind the genera *Diaporthe* (*Sordariomycetidae*; 27.7 ± 2.09%), *Xanthoria* (OSLEUM clade; 25.9 ± 2.14%) and members belonging to the enriched order *Xylariales* (24.08 ± 3.01%) with contributions from densely populated genera *Hypoxylon* (24.72 ± 2.69%), *Annulohypoxylon* (25.16 ± 1.73%), *Nemania* (25.44 ± 2.92%), *Neopestalotiopsis* (27.5 ± 1.08%), *Pestalotiopsis* (27.39 ± 1.56%), *Arthrinium* (26.12 ± 3.34%), *Apiospora* (26.42 ± 2.87%) and *Hymenoscyphus* (24.66 ± 2.11%). Sister taxon *Scedosporium* (24.59 ± 1.66%) exhibited slightly lower AA density and belonged above the 90^th^ percentile. Interestingly, members of the *Taphrinomycotina* (7.53 ± 3.36%) and *Saccharomycetales* (8.95 ± 4.63%), often associated with lignocellulose deconstruction, displayed significantly reduced AA abundance in stark contrast to neighbouring phylogenies such as *Pezizomycotina* (18.42 ± 4.29%). Members of the order *Helotiales* (21.04 ± 3.42%) and class *Dothideomycetes* (20.26 ± 3.98%) displayed a degree of enrichment of AAs whilst members belonging to *Eurotiomycetes* (16.64 ± 2.83%) displayed lower abundances. Considering how the AA class of enzymes is predominantly associated with the degradation of lignin and crystalline cellulose it highlights a potential strategy of the fungus to target these components. Indeed, in a mixed microbial community grown on wheat straw the fungus was observed to become more dominant in the later stages of the culture, potentially due to its capacity to modify the more difficult to degrade components of lignocellulose for growth (5).

**Figure 2.**
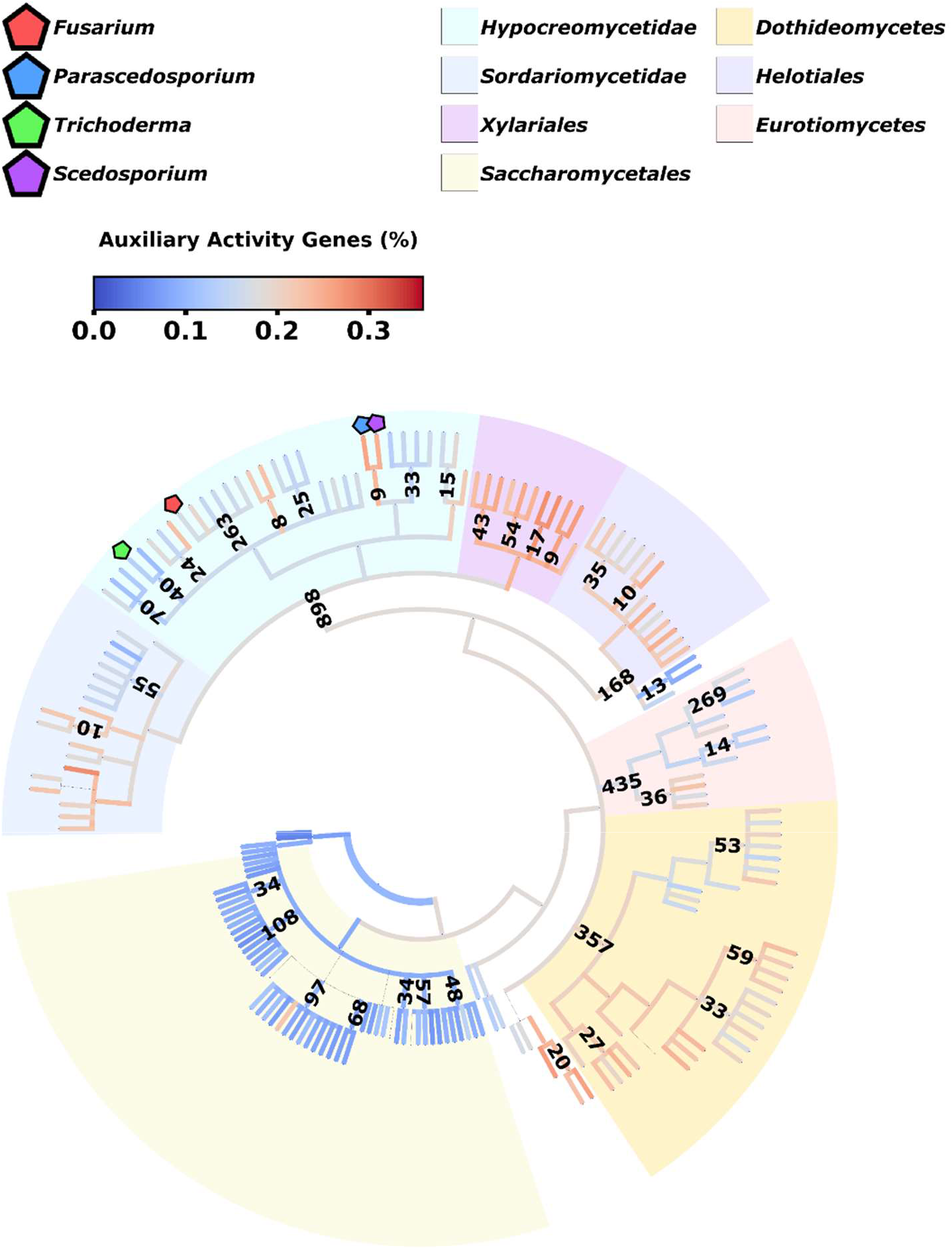
Auxiliary activity distribution and density across the ascomycete tree of life. Genes predicted for ascomycete genome assemblies were annotated for CAZymes to explore patterns in the distribution and density of auxiliary activities (n=2570) within the ascomycete phylogenetic tree. Branch colors indicate the mean proportion of auxiliary activities within all CAZyme annotations accounted for by all descendant taxa. Yellow bubbles and annotations represent number of sequenced genomes available. Key taxa have been highlighted. Genera and families with less than 3 and 8 species level representatives, respectively, have been pruned for clarity (n=462 taxa). Nodes of taxonomic ranks below genus have been pruned (n=93 taxa).

Within white- and brown- (basidiomycete), and soft-rot (ascomycete) fungi, it has been demonstrated that the CAZyme repertoire can vary greatly from species to species (8). To investigate the repertoire of *P. putredinis* NO1 in more detail, CAZyme domains were compared to that of three other wood-degrading ascomycetes. *Scedosporium boydii* is located within the sister taxon of *Parascedosporium* and has a genome of 43 Mb containing 1029 CAZyme domains. The genome and CAZyme complement of the soft-rot *P. putredinis* NO1 are larger than that of *Trichoderma reesei* which contains 786 domains in 34 Mb of DNA. *T. reesei* is a mesophilic soft-rot fungus known for its ability to produce high titres of polysaccharide-degrading enzymes that are used in biomass-degrading enzyme cocktails (9). The genome of *P. putredinis* NO1 is slightly smaller than that of *Fusarium oxysporum* at 47 Mb, a phytopathogenic fungus containing an expanded CAZyme repertoire of 1430 domains (10). The lignocellulose degrading activities of *F. oxysporum* have been well-investigated in part due to its pathogenicity and ability to ferment sugars from lignocellulose breakdown directly into ethanol (11, 12).

Examining the distribution of predicted CAZyme domains revealed that despite the similar overall number of CAZyme domains for *P. putredinis* NO1 and *T. reesei*, the proportion of AA class CAZyme domains is much higher in the genome of *P. putredinis* NO1 (**Figure 1A**). Proportionally, AA class CAZymes make the largest contribution to CAZyme repertoire in the genome of *P. putredinis* NO1 compared to the other ascomycetes (**Figure 1**). This again could suggest an oxidative strategy to target to lignin and crystalline cellulose. Although analysis of fungal secretomes would be required to confirm an improved ability of *P. putredinis* NO1 to deconstruct lignocellulosic components, the high potential capacity for degradation of lignin and crystalline cellulose within the genome suggests that this is an important fungus to explore for new lignocellulose-degrading enzymes. Especially considering that this is the first genome assembly of the genus *Parascedosporium*.

The increased contribution of AA class CAZymes is mirrored by a reduced proportion of GH class CAZymes in the *P. putredinis* NO1 genome compared to *T. reesei* and *F. oxysporum*. This reduced GH contribution is also visible in the genome of *S. boydii*, a close relative of *P. putredinis* NO1. Despite the reduced number of the hydrolytic GH class CAZymes, the repertoires of *P. putredinis* NO1 and *S. boydii* contain the highest proportions of CBMs, domains typically associated with hydrolytic CAZymes such as GHs (13), but which have also been observed in oxidative LPMOs (14, 15). The increased proportion of CBMs in the genome of *P. putredinis* NO1 could aid the catalytic CAZymes in accessing and binding to these substrates. Indeed, examining the CBM domains at the family level shows a high number of crystalline cellulose binding domains (CBM1) in the genome of both *P. putredinis* NO1 and *S. boydii*, much higher than the number of domains assigned to any of the other CBM families (**Supplementary** Figure 1).

### Closer Investigation of the AA CAZyme Repertoire Reveals More About the Lignocellulose Degrading Strategy of *P. putredinis* NO1

The high number of AA domains, a functional class that notably contains LPMOs, peroxidases, and laccases (2), in the genome of *P. putredinis* NO1 are likely to endow this fungus with the ability to degrade recalcitrant components of the plant cell wall through a primarily oxidative mechanism. LPMOs are copper-containing enzymes that enhance polysaccharide degradation by generating new sites for attack by hydrolytic CAZymes (16). LPMOs have been shown to act on all major polysaccharide components of lignocellulose. Their oxidative action relies on exogenous electron donors provided by other AA family CAZymes, small molecule reductants and even lignin (2, 16). It has recently been demonstrated that LPMOs readily utilise hydrogen peroxide (H2O2) as a cosubstrate also (17, 18).

Investigating the distribution of AA domains across the AA families revealed AA9 family members to be the most abundant in the *P. putredinis* NO1 genome with 35 domains, the highest in the four ascomycetes investigated here (**Supplementary** Figure 2). This family contains the cellulose, xylan, and glucan-active LPMOs described above (19). AA3 and AA3_2 domains are the second and third most abundant families in the *P. putredinis* NO1 genome with 29 and 27 domains, respectively. These are flavoproteins of the Glucose-methanol-choline (GMC) oxidoreductase family which includes activities such as cellobiose dehydrogenase, glucose-1-oxidase, aryl alcohol oxidase, alcohol oxidase and pyranose oxidase (20). It is proposed that flavin binding oxidative enzymes of this family play a central role in spatially and temporally supplying H2O2 to LPMOs and peroxidases or to produce radicals that degrade lignocellulose through Fenton chemistry (17). The *P. putredinis* NO1 genome also contains 12 AA7 family domains, the family of glucooligosaccharide oxidase enzymes. These have recently been demonstrated to transfer electrons to AA9 LPMOs which boosts cellulose degradation (21). Altogether, the apparent expansion of these LPMO system families suggest a potentially increased capacity for *P. putredinis* NO1 to oxidatively target crystalline cellulose.

The genome of *P. putredinis* NO1 also contains 12 AA1 family CAZyme domains. This family includes laccase and multi-copper oxidase enzymes which catalyse the oxidation of various aromatic substrates while simultaneously reducing oxygen to water (22). It has also been demonstrated that laccases can boost LPMO activity through the release of low molecular weight lignin polymers from biomass which can in turn donate electrons to LPMOs (23). Additionally, 7 domains belonging to the AA8 family were identified, a family of iron reductase domains initially identified as the N-terminal domain in cellobiose dehydrogenase enzymes but also found independently and appended to CBMs (2, 24, 25). These domains are believed to be involved in the generation of reactive hydroxyl radicals that can indirectly depolymerize lignin. There are 6 AA4 domains in the genome of *P. putredinis* NO1, the highest number of the four ascomycetes investigated here. These are vanillyl-alcohol oxidase enzymes with the ability to catalyse the conversion of a wide range of phenolic oligomeric compounds (26). These may act downstream of the lignin depolymerisation catalysed by other members of the AA class. There is a clear capacity in the *P. putredinis* NO1 genome for lignin depolymerisation and metabolism through the multiple domains identified belonging to these families. The *P. putredinis* NO1 genome also contains two AA16 domains, a recently identified family of LPMO proteins with an atypical product profile compared to the traditional AA9 family LPMOs and a potentially different mode of activation (27).

Gene expression of CAZymes in the *P. putredinis* NO1 genome has been explored previously during growth on glucose, compared to growth on wheat straw with samples taken at days 2, 4, and 10 (5). This transcriptomic data gives a view of the potential strategy by which *P. putredinis* NO1 utilises its expanded repertoire of AA class CAZymes. Up-regulation of AA class CAZymes during growth on wheat straw compared to growth on glucose was observed predominantly at day 4 and then gave way to up-regulation instead of mainly GH class hydrolytic CAZymes at day 10. This could represent a strategy where the recalcitrant lignin and crystalline cellulose are targeted first by LPMOs and lignin degraders such as laccases, making the polysaccharide substrates of hydrolytic GH enzymes more accessible.

### Searching the *P. putredinis* NO1 Genome for New Oxidative Lignocellulose Degrading Enzymes with Sequence-, Domain-, and Structural-Based Strategies

Due to the evidence of a strategy for *P. putredinis* NO1 to target the most recalcitrant components of lignocellulose and the recent discovery of a new oxidase with the ability to cleave the major linkage in lignin from this strain (5), it was hypothesised the genome of this fungus contains additional new enzymes for the breakdown of plant biomass. Particularly this fungus could contain new enzymes with roles in degrading the lignin and crystalline cellulose components and which have not been annotated as CAZymes in this analysis.

Traditionally, homologue searching has been performed using a sequence-based approach (28). Using either the primary amino acid sequence of an example protein to search an unknown database for similar sequences, or with the use of Hidden Markov Models (HMMs) to search for domains of interest (29). However, both techniques rely on primary amino acid sequence homology and neglect that proteins with distantly related sequences may have similar three-dimensional structures and therefore activity. The recent emergence of AlphaFold provides a resource for the fast and accurate prediction of unknown protein structures (30). Using this tool, structures were predicted for >96% of the protein-coding regions of the *P. putredinis* NO1 genome. These structures were used to create a database of protein structures into which structures of interesting enzymes such as those for LPMOs, laccases, and peroxidases could be searched. These structural searches for new enzymes were performed alongside sequence- and domain-based searches for comparison of the ability to identify interesting new candidates.

LPMO related sequences were searched for in the *P. putredinis* NO1 genome using the sequence of an AA9 family LPMO from *Aspergillus niger* with the default E-value cut off of 1 x 10^-5^, with the AA9 HMM from Pfam and considering domain hits that fell within the default significance inclusion threshold of 0.01 (31), and the structure of the same *A. niger* LPMO with a tailored ‘lowest percentage match’ parameter. In total, 49 sequences were identified across the three searching strategies and 33 of these sequences were also annotated by dbCAN as AA9 family LPMOs (**Supplementary Table 1**). With the objective of identifying new enzymes, the remaining 16 sequences were investigated further, and the distribution of the identification of these sequences across the three search strategies can be seen in **Table 1**.Two of the sequences, PutMoI and PutMoM, were identified by all three search approaches. These sequences both had conserved signal peptides with a conserved N-terminal histidine after the cleavage site, a characteristic feature of LPMOs (32).

**Table 1.** Identifying LPMO related proteins encoded in the *P. putredinis* NO1 genome. Coding regions of proteins related to LPMOs identified through genome searching approaches with the sequence of an *A. niger* AA9 LPMO (E-value cut-off = 1 x 10^-5^), the Pfam AA9 HMM (Significance threshold = 0.01), and the structure of the *A. niger* AA9 LPMO (Lowest percentage match = 50%) and which were not annotated as AA9 CAZymes by dbCAN. InterPro annotations were retrieved where possible.

| Coding Region | GenBank Accession | Protein ID | Identified by Searching Approach |  |  | Interpro Annotation |
| --- | --- | --- | --- | --- | --- | --- |
|  |  |  | Sequence | Domain | Structure |  |
| FUN_000653-T1 | CAI7987917.1 | PutMoA |  |  | ✓ | AA16 LPMO |
| FUN_000713-T1 | CAI7987978.1 | PutMoB |  |  | ✓ | Rho factor associated |
| FUN_002573-T1 | CAI7991617.1 | PutMoC |  | ✓ |  | - |
| FUN_002890-T1 | CAI7992277.1 | PutMoD |  |  | ✓ | AA16 LPMO |
| FUN_002962-T1 | CAI7992399.1 | PutMoE |  |  | ✓ | - |
| FUN_003190-T1 | CAI7992922.1 | PutMoF |  |  | ✓ | Ferritin-like |
| FUN_003535-T1 | CAI7993628.1 | PutMoG |  | ✓ |  | AA13 LPMO |
| FUN_003783-T1 | CAI7994168.1 | PutMoH |  |  | ✓ | - |
| FUN_006366-T1 | CAI7999797.1 | PutMoI | ✓ | ✓ | ✓ | AA9 LPMO |
| FUN_006413-T1 | CAI7999893.1 | PutMoJ |  | ✓ |  | AA9 LPMO |
| FUN_006553-T1 | CAI8000144.1 | PutMoK |  |  | ✓ | - |
| FUN_007242-T1 | CAI8001774.1 | PutMoL |  | ✓ |  | AA9 LPMO |
| FUN_007666-T1 | CAI8002525.1 | PutMoM | ✓ | ✓ | ✓ | AA9 LPMO |
| FUN_008106-T1 | CAI8003467.1 | PutMoN | ✓ | ✓ |  | AA9 LPMO |
| FUN_009239-T1 | CAI7992001.1 | PutMoO |  |  | ✓ | - |
| FUN_010012-T1 | CAI8003342.1 | PutMoP |  |  | ✓ | - |

When creating the structure database it was tempting to filter predicted structures by pLDDT score, the AlphaFold metric for prediction confidence, to create a database solely of ‘high confidence’ structures (30). However, pLDDT scores reflect local confidence and should instead be used for assessment of individual domains (33). The majority of the structures generated here had pLDDT score of over 60%, however pLDDT scores lower than 70% are considered low confidence (**Supplementary** Figure 3). Extracellular enzymes are of particular interest here, but these often have disordered N-terminal signal peptides which can reduce the overall pLDDT scores. Therefore, for secreted enzyme identification from AlphaFold structures it is inappropriate to filter by pLDDT score. Indeed, the PutMoI structure mentioned above had a pLDDT score of 62%, considered to be low confidence (30), but which had characteristic features of LPMOs and which demonstrated structural similarity to the *A. niger* AA9 LPMO used for structural searches (**Figure 3A and 3B**). The central beta-sheet structures align well to the *A. niger* AA9 LPMO for both PutMoI and PutMoM, but both also have additional loops of disordered protein which likely explains the relatively low PDBefold alignment confidence scores (Q-scores) of 0.23 and 0.34 for PutMotI and PutMoM, respectively. This again highlights the unreliability of structural confidence scores alone and demonstrates how manual inspection of structural alignments may prove more useful. Despite not being annotated as AA9 LPMOs by the dbCAN server for CAZyme annotation (7), both sequences were identified using the Pfam AA9 HMM and appear to be conserved AA9 LPMOs and, therefore, are not of interest in the discovery of new enzymes.

**Figure 3.**
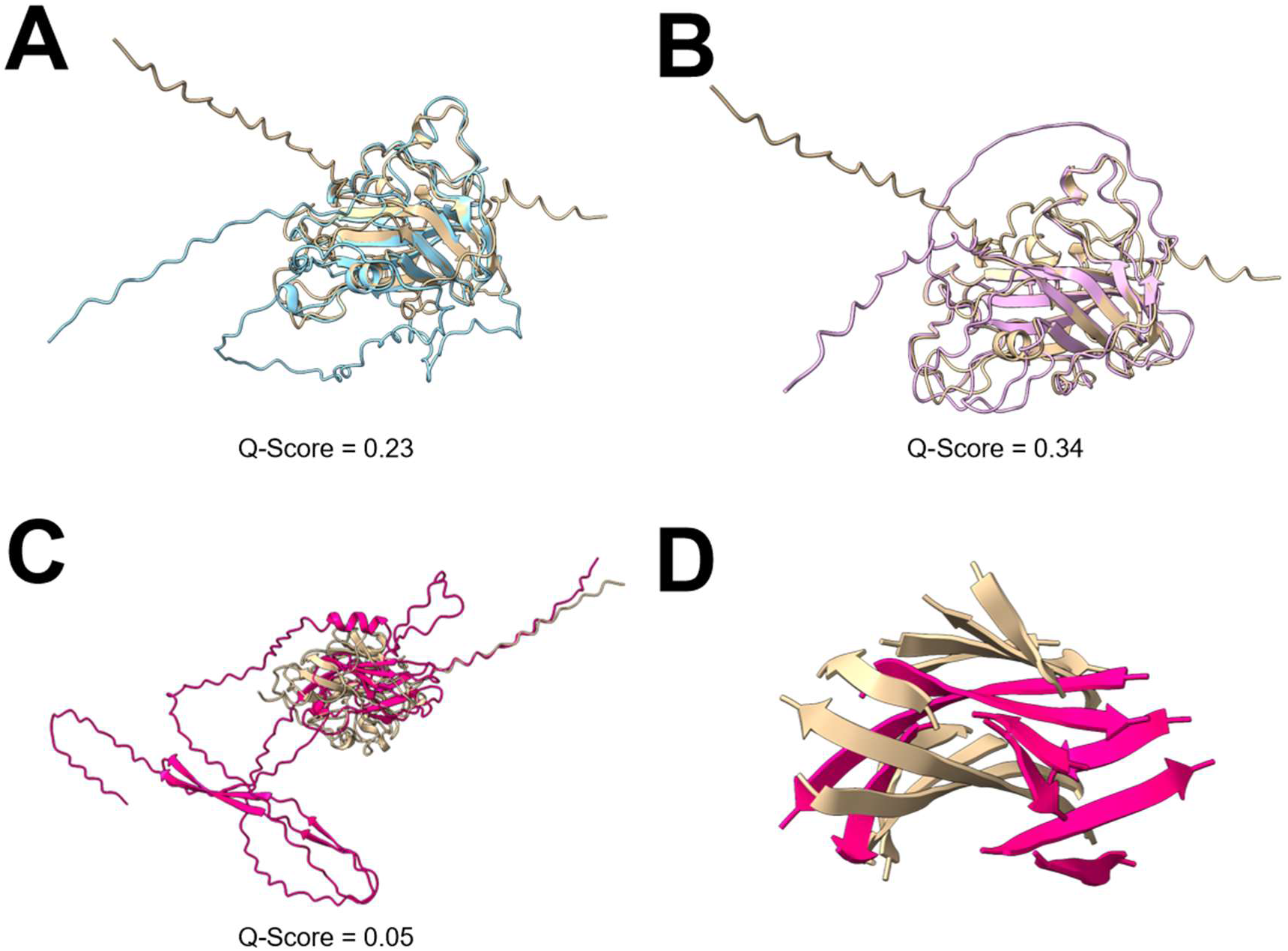
Structural comparison of LPMO related proteins. The AlphaFold predicted structures of three sequences, PutMoI (**A**), PutMoM (**B**), and PutMoP (**C** and **D**) from the *P. putredinis* NO1 genome structurally aligned to the *A. niger* AA9 LPMO used in sequence and structure-based searching (UniProt ID: A2QZE1). *A. niger* AA9 LPMO (Beige), PutMoI (Blue), PutMoM (Pink), PutMoP (Hot Pink). Q-score is a quality function of Cα alignment from PDBefold.

By utilising multiple searching approaches, potentially new sequences with LPMO related activities can be identified. When searching for LPMO related sequences, domain-based approaches identified all coding regions also identified by sequence-based searching as well as additional coding regions (**Supplementary Table 1**). This pattern of domain-based searching identifying more coding regions than sequence-based searching was also observed for the other activities investigated (**Supplementary Tables 2 and 3**). For structure-based searching, parameters of the searches could be tailored to identify additional coding regions with lower overall structural similarity, but which may still be interesting. For example, searching the against the *P. putredinis* NO1 genome structure database with the structure of the *A. niger* AA9 LPMO, and with the ‘lowest acceptable match’ parameter which is the cutoff at which secondary structures must overlap between a query and a target set at 50 %, yielded 30 coding regions (**Supplementary Table 1**). Of these sequences, 9 were not identified by the sequence or domain-based searching approaches and were investigated in more detail (**Table 1**). To investigate these further, sequences were searched against the NCBI non- redundant protein database to identify related sequences (34), conserved domains were predicted with InterPro any CAZyme domains were annotated with dbCAN (7), the predicted structures were compared with structures in the PDB database (35), and secretion signal peptides were predicted with SignalP (36) in an attempt to elucidate the potential functions. Two of the sequences, PutMoA and PutMoD, are the two predicted AA16 LPMOs identified in the *P. putredinis* NO1 CAZyme repertoire earlier (**Supplementary** Figure 2). Another two sequences, PutMoH and PutMoK, were not annotated as CAZymes but had conserved BIM1- like domains. BIM1-like proteins are LPMO_auxilliary-like proteins, function in fungal copper homeostasis, and share a similar copper coordination method to the LPMOs which they are related to (37). Although not likely to be involved in lignocellulose breakdown, this highlights how structurally related proteins in terms of active site or co-factor coordination structures can be identified with structural approaches where sequence- and domain-based approaches fail. Three of the nine sequences were also identified as being upregulated when *P. putredinis* NO1 was previously grown on wheat straw and compared to growth on glucose (**Supplementary File 1**) (5). Although this does not confirm the role of these proteins in lignocellulose breakdown, it provided another layer of information for the selection of interesting candidate sequences to investigate further. PutMoP was the most interesting sequence identified solely by the structural searching and showing upregulation during growth on wheat straw compared to glucose. It was not annotated as a CAZyme, no conserved domains were identified, and sequence homology was only observed to hypothetical proteins in the NCBI non-redundant protein database (34). Comparing the AlphaFold predicted structure of PutMoP to the *A. niger* AA9 LPMO revealed similarity at the central beta-sheet structure despite a very low Q-score of 0.05 (**Figure 3C and 3D**). A secretion signal peptide was also predicted for this protein, suggesting an extracellular role. This immunoglobulin-like distorted β-sandwich fold is a characteristic structural feature of LPMOs and is shared across the LPMO CAZyme families (38). The similarity of this central structure is likely the reason for identification of this sequence by structural comparison. This structural similarity at the protein centre, the lack of amino acid sequence similarity, and the conserved secretion signal makes this protein an interesting candidate for further investigation. Searching the PutMoP structure against the whole PDB structure database returned many diverse proteins not linked to lignocellulose breakdown, however the Q-score was very low for all the structures and did not help to discern the potential activity of this protein. The sequence lacks the N-terminal histidine after the signal peptide cleavage site which is conserved in LPMOs so this protein is unlikely to be an LPMO. However, a secreted unknown protein with some central structural similarity to an important class of oxidative proteins that degrade crystalline cellulose is of definite interest.

In addition to searching for LPMO related sequences, classes of enzymes involved in the breakdown of lignin are important targets for the biorefining of plant biomass. The recalcitrance of lignin is a limiting factor hindering the industrial use of lignocellulose as a feedstock to produce biofuels. Lignin itself is also a historically underutilised feedstock for valuable chemicals (39). Laccases are multicopper oxidase family enzymes that catalyse oxidation of phenolic compounds through an electron transfer reaction that simultaneously reduces molecular oxygen to water (23). They modify lignin by depolymerisation and repolymerisation, Cα oxidation, and demethylation and are particularly efficient due to their use of readily available molecular oxygen as the final electron acceptor (40, 41).

Laccase related seqeunces were searched for in the *P. putredinis* NO1 genome using the sequence of an AA1 family laccase from *A. niger*, a bespoke HMM constructed from ascomycete laccase and basidiomycete multi-copper oxidase sequences downloaded from the laccase engineering database (42), and with the structure of the *A. niger* AA1 laccase. In total, 32 sequences were identified across the three searching strategies and only 9 of these were annotated by dbCAN as AA1 family CAZymes (**Supplementary Table 2**). The bespoke HMM allowed for more divergent sequences for these enzymes to be incorporated into the model’s construction. The result was the identification of sequences that when explored further looked like laccase enzymes but were missed by traditional CAZyme annotation, highlighting how searching for CAZymes alone is a limited method for identifying lignocellulose degrading enzymes. However, for the identification of new lignocellulose degrading enzymes, more divergent sequences are of interest. A single coding sequence, PutLacJ was identified by the structural searching approach with a 30% ‘lowest acceptable match’ parameter that was not identified by sequence or domain-based searching (**Table 2**).

**Table 2.**
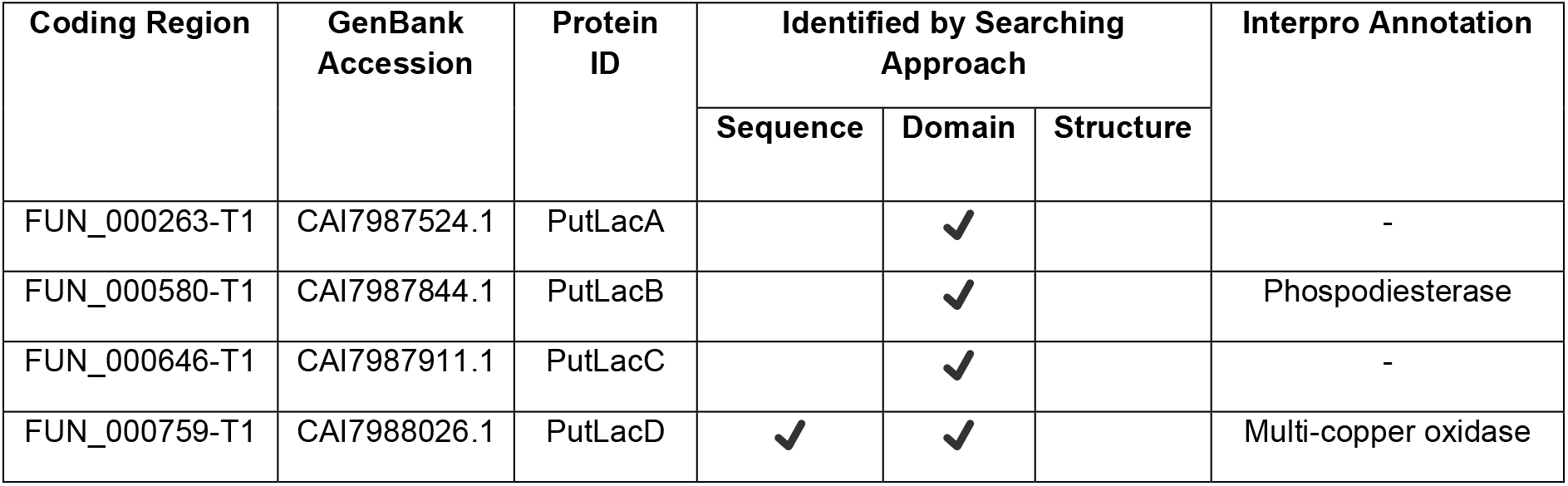
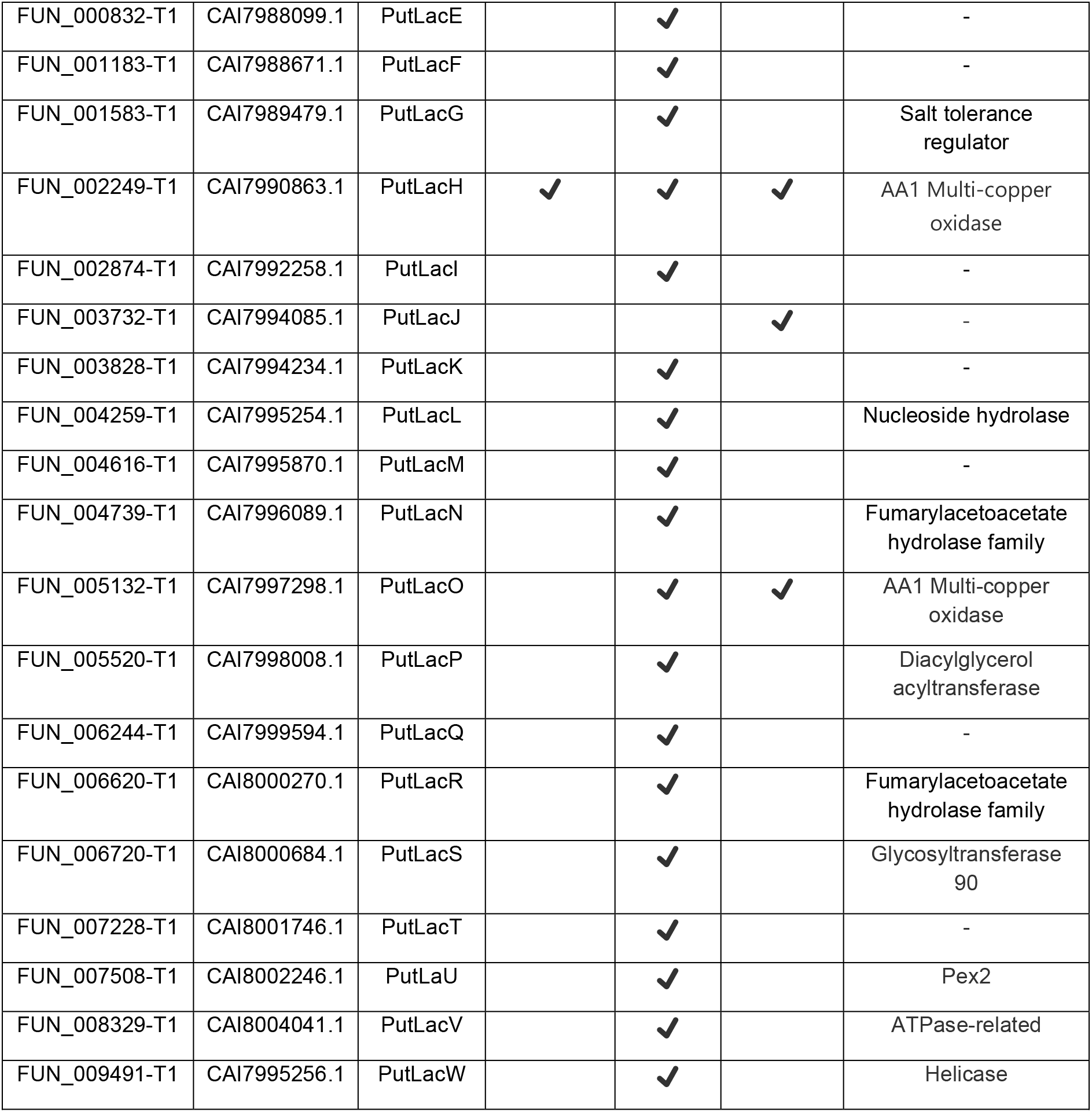
Identifying laccase related proteins encoded in the *P. putredinis* NO1 genome. Coding regions of proteins related to laccases identified through genome searching approaches with the sequence of an *A. niger* AA1 laccase (E-value cut-off = 1 x 10^-5^), the bespoke laccase and multicopper oxidase HMM constructed from sequences from the laccase engineering database (Significance threshold = 0.01), and the structure of the *A. niger* AA1 laccase (Lowest percentage match = 30%) and which were not annotated as AA1 CAZymes by dbCAN. InterPro annotations were retrieved where possible.

| Coding Region | GenBank Accession | Protein ID | Identified by Searching Approach |  |  | Interpro Annotation |
| --- | --- | --- | --- | --- | --- | --- |
|  |  |  | Sequence | Domain | Structure |  |
| FUN_000263-T1 | CAI7987524.1 | PutLacA |  | ✓ |  | - |
| FUN_000580-T1 | CAI7987844.1 | PutLacB |  | ✓ |  | Phosphodiesterase |
| FUN_000646-T1 | CAI7987911.1 | PutLacC |  | ✓ |  | - |
| FUN_000759-T1 | CAI7988026.1 | PutLacD | ✓ | ✓ |  | Multi-copper oxidase |
| FUN_000832-T1 | CAI7988099.1 | PutLacE |  | ✓ |  | - |
| FUN_001183-T1 | CAI7988671.1 | PutLacF |  | ✓ |  | - |
| FUN_001583-T1 | CAI7989479.1 | PutLacG |  | ✓ |  | Salt tolerance regulator |
| FUN_002249-T1 | CAI7990863.1 | PutLacH | ✓ | ✓ | ✓ | AA1 Multi-copper oxidase |
| FUN_002874-T1 | CAI7992258.1 | PutLacI |  | ✓ |  | - |
| FUN_003732-T1 | CAI7994085.1 | PutLacJ |  |  | ✓ | - |
| FUN_003828-T1 | CAI7994234.1 | PutLacK |  | ✓ |  | - |
| FUN_004259-T1 | CAI7995254.1 | PutLacL |  | ✓ |  | Nucleoside hydrolase |
| FUN_004616-T1 | CAI7995870.1 | PutLacM |  | ✓ |  | - |
| FUN_004739-T1 | CAI7996089.1 | PutLacN |  | ✓ |  | Fumarylacetoacetate hydrolase family |
| FUN_005132-T1 | CAI7997298.1 | PutLacO |  | ✓ | ✓ | AA1 Multi-copper oxidase |
| FUN_005520-T1 | CAI7998008.1 | PutLacP |  | ✓ |  | Diacylglycerol acyltransferase |
| FUN_006244-T1 | CAI7999594.1 | PutLacQ |  | ✓ |  | - |
| FUN_006620-T1 | CAI8000270.1 | PutLacR |  | ✓ |  | Fumarylacetoacetate hydrolase family |
| FUN_006720-T1 | CAI8000684.1 | PutLacS |  | ✓ |  | Glycosyltransferase 90 |
| FUN_007228-T1 | CAI8001746.1 | PutLacT |  | ✓ |  | - |
| FUN_007508-T1 | CAI8002246.1 | PutLaU |  | ✓ |  | Pex2 |
| FUN_008329-T1 | CAI8004041.1 | PutLacV |  | ✓ |  | ATPase-related |
| FUN_009491-T1 | CAI7995256.1 | PutLacW |  | ✓ |  | Helicase |

PutLacJ was not annotated as a CAZyme by dbCAN but does have a predicted cupredoxin domain, a feature of laccase enzymes (43). Structural comparisons against the PDB structure database revealed alignments with moderate confidence scores to copper-containing nitrite reductases from *Neisseria gonorrhoeae* which are suggested to play a role in pathogenesis (44). In fungi, it is more likely that these are playing a role in denitrification (45). The lack of a signal peptide make it unlikely that this protein is involved in lignin depolymerisation, despite the structural similarity to the beta-sheet regions of the *A. niger* laccase (**Figure 4**).

**Figure 4.**
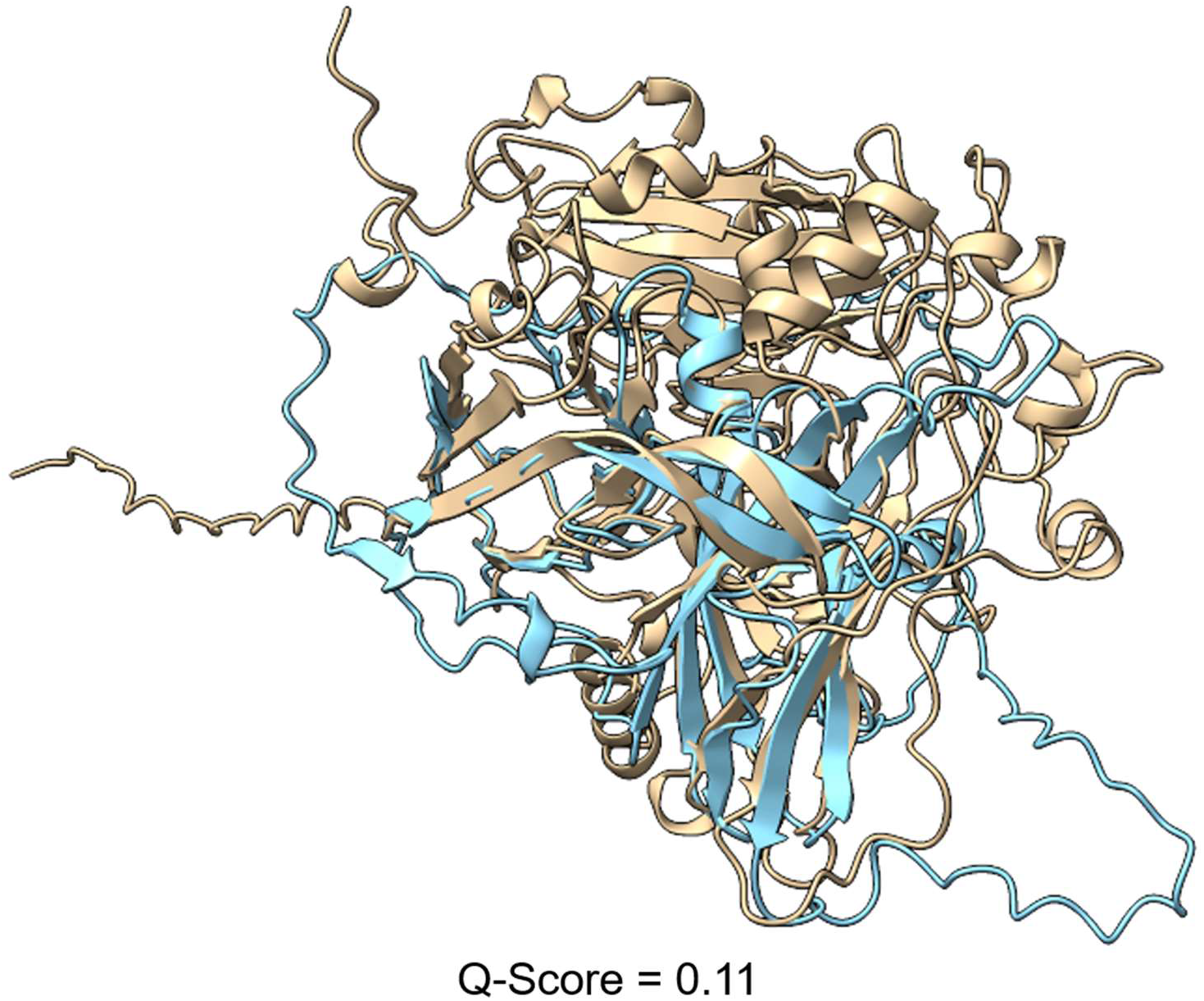
Structural comparison of PutLacJ laccase related protein. The AlphaFold predicted structures of the sequence PutLacJ from the *P. putredinis* NO1 genome structurally aligned to the *A. niger* laccase used in sequence and structure-based searching (UniProt ID: A2QB28). *A. niger* laccase (Beige), PutLacJ (Blue). Q-score is a quality function of Cα alignment from PDBefold.

Peroxidases (PODs) also play a major role in lignin deconstruction by white-rot fungi. PODs are lacking in brown-rot species, presumably due to their non-ligninolytic specialisation of substrate degradation (46). The identification of new putative peroxidases in *P. putredinis* NO1 is of interest. Fungal class II peroxidases are divided into three lignolytic forms; lignin peroxidase (LiP), manganese peroxidase (MnP), and versatile peroxidase (VP) (47).

Sequence searches into the *P. putredinis* NO1 genome using sequences of MnP from *Aureobasidium subglaciale*, LiP from *F. oxysporum*, and VP from *Pyronema confluens* only yielded 2 sequences (**Supplementary Table 3**). Both peroxidase related sequences were also identified by domain searching using a bespoke HMM constructed from sequences of MnPs, LiPs, and VPs downloaded from the fPoxDB database of peroxidase sequences (48). This domain-based approach only identified 3 sequences in total, all of which were annotated as AA2 family CAZymes also (**Supplementary Table 3**). However, structural-based searching using the structures of the same three peroxidases, and with a ‘lowest acceptable match’ parameter of 30% used in sequence-based searches identified 9 coding regions in total (**Supplementary Table 3**), 7 of which were not identified by sequence- or domain-based searching approaches and were not annotated as AA2 CAZymes (**Table 3**), but were all found to be upregulated previously when *P. putredinis* NO1 was grown on wheat straw compared to growth on glucose (**Supplementary File 1**) (5).

**Table 3.** Identifying peroxidase related proteins encoded in the *P. putredinis* NO1 genome. Coding regions of proteins related to peroxidases identified through genome searching approaches with the sequences of an MnP from *A. subglaciale*, LiP from *F. oxysporum*, and VP from *P. confluens* (E-value cut-off = 1 x 10^-5^), the bespoke peroxidase HMM contricted from MnP, LiP, and VP seqeunces in the fPoxDB database (Significance threshold = 0.01), and the structure of the same three peroxidases used for seqeunce searches (Lowest percentage match = 30%) and which were not annotated as AA2 CAZymes by dbCAN. InterPro annotations were retrieved where possible.

| Coding Region | GenBank Accession | Protein ID | Identified by Searching Approach |  |  | Interpro Annotation |
| --- | --- | --- | --- | --- | --- | --- |
|  |  |  | Sequence | Domain | Structure |  |
| FUN_002995-T1 | CAI7992466.1 | PutPoxA |  |  | ✓ | DUF3632 |
| FUN_003542-T1 | CAI7993642.1 | PutPoxB |  |  | ✓ | Arabinofuranosidase |
| FUN_003618-T1 | CAI7993895.1 | PutPoxC |  |  | ✓ | - |
| FUN_004484-T1 | CAI7995643.1 | PutPoxD |  |  | ✓ | Cell division control |
| FUN_008413-T1 | CAI8004205.1 | PutPoxE |  |  | ✓ | SIT4 phosphatase-associated |
| FUN_008923-T1 | CAI7988420.1 | PutPoxF |  |  | ✓ | - |
| FUN_009329-T1 | CAI7993214.1 | PutPoxG |  |  | ✓ | - |

Investigating these sequences further revealed two sequences to be the most interesting, PutPoxA and PutPoxG, both with low Q-scores of 0.01 and 0.04, respectively. PutPoxA was not annotated as a CAZyme but does have a predicted domain of unknown function family 3632 (DUF3632). Genes encoding DUF3632 domains were previously found to be upregulated in the filamentous ascomycete *Neurospora crassa* when the CLR-2 transcription factor, important for growth on cellulose, was constitutively expressed (49). The protein does however lack a signal peptide and structural comparison to the *A. subglaciale* MnP shows similar helical structures, but these secondary structures do not appear to overlap very well (**Figure 5A**). PutPoxG was not annotated as a CAZyme and no conserved domains were identified, although the helical structures do seem to align better with the *A. subglaciale* MnP than PutPoxA (**Figure 5B**). Furthermore, searching of both structures against the PDB database was performed, but all alignments had very low Q-scores of less than 0.1.

**Figure 5.**
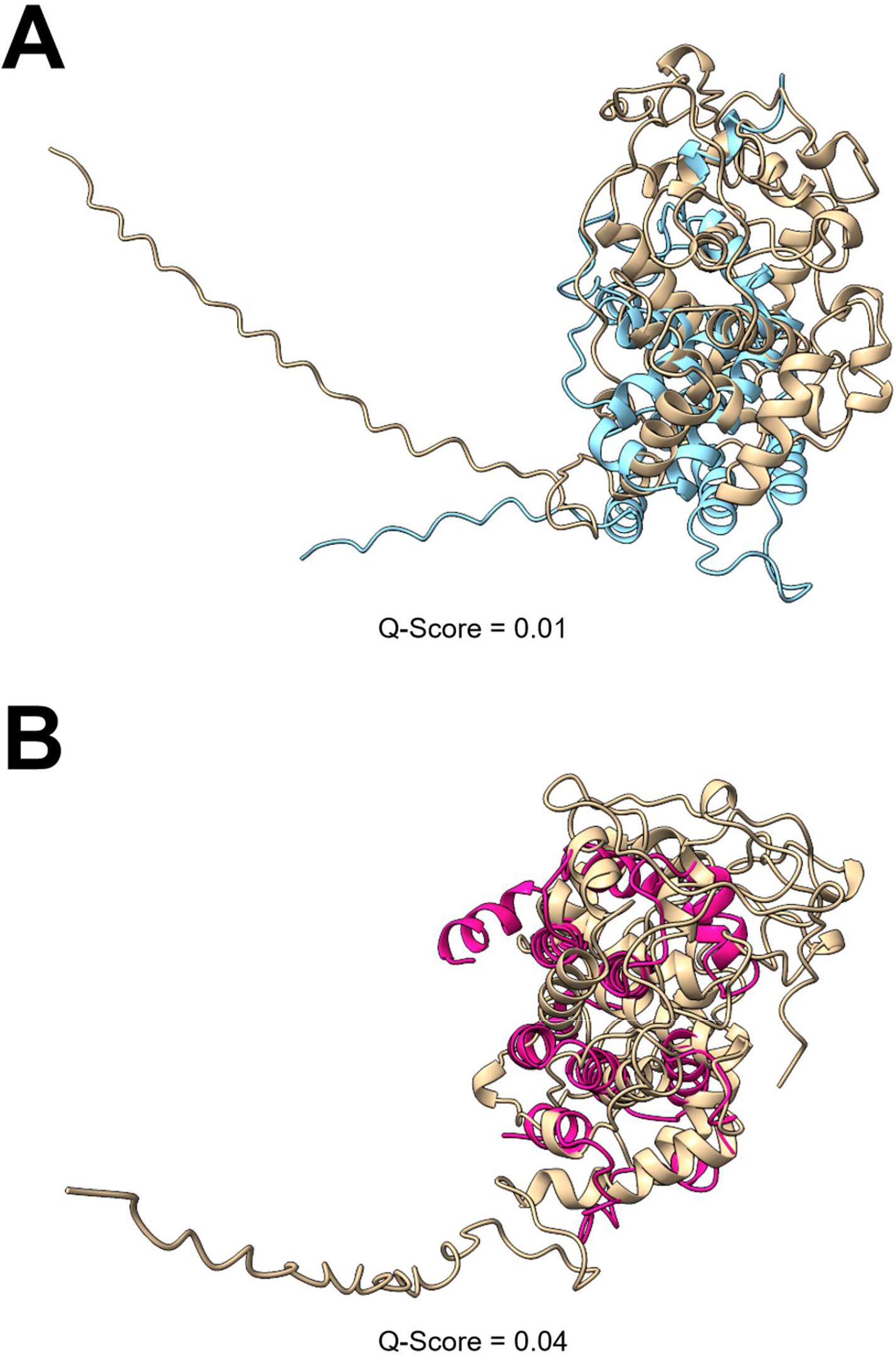
Structural comparison of peroxidase related proteins. The AlphaFold predicted structures of two sequences, PutPoxA (**A**) and PutPoxG (**B**), from the *P. putredinis* NO1 genome structurally aligned to the *A. subglaciale* MnP used in sequence and structure-based searching. A predicted structure was unavailable and so a predicted structure was generated with AlphaFold. *A. subglaciale* MnP (Beige), PutPoxA (Blue), PutPoxG (Hot Pink). Q-score is a quality function of Cα alignment from PDBefold.

As with the candidates identified by LPMO and laccase searching approaches, it is hard to be confident on sequence and structural investigation alone that these proteins are involved in lignocellulose breakdown. Although by utilising multiple searching approaches, more divergent and varied sequences with potential relation to industrially important enzymes have been identified here. This strategy of searching for new enzymes involved in the breakdown of the most recalcitrant components of lignocellulose would work well when combined with additional layers of biological data e.g., transcriptomic, or proteomic data. Many of the coding regions investigated here show structural similarity to the interesting classes of enzymes with which they were identified but lack the sequence similarity and therefore the functional annotation. Transcriptomic data showing upregulation of these genes or proteomic data showing increased abundances of these proteins when the organism in question is grown on lignocellulosic substrates would inspire more confidence in the role of these proteins in the degradation of plant-biomass. Therefore, we used sequence similarity to identify the corresponding transcripts for these coding regions in the transcriptomic time course dataset of *P. putredinis* NO1 grown for 10 days in cultures containing wheat straw published previously (5). The transcriptomic data was explored for all sequences which were identified solely by structural searches and therefore considered interesting (**Supplementary File** 1). For the four sequences explored in more detail, we found that three of the four: PutMoP, PutPoxA, and PutPoxG, were found to be significantly upregulated on at least one timepoint when grown on wheat straw compared to growth on glucose (**Figure 6**). The remaining sequence, PutLacJ, expression was found to be significantly higher during growth on glucose compared to growth on wheat straw. However structural investigation revealed that PutLacJ had similarity to copper-containing nitrite reductase proteins and it was concluded that it is unlikely to be involved in lignocellulose breakdown. Characterisation would be required to confirm the role of these candidates in lignocellulose breakdown and to understand whether these activities are new. However, the implication in lignocellulose degrading processes through the analysis of transcriptomic data provides another source of information by which candidates identified through the described strategy can be investigated. It is hoped that adoption of a similar strategy for analysis of the wealth of sequence data now publicly available will allow identification of novel enzyme sequences for many important processes to be made simpler.

**Figure 6.**
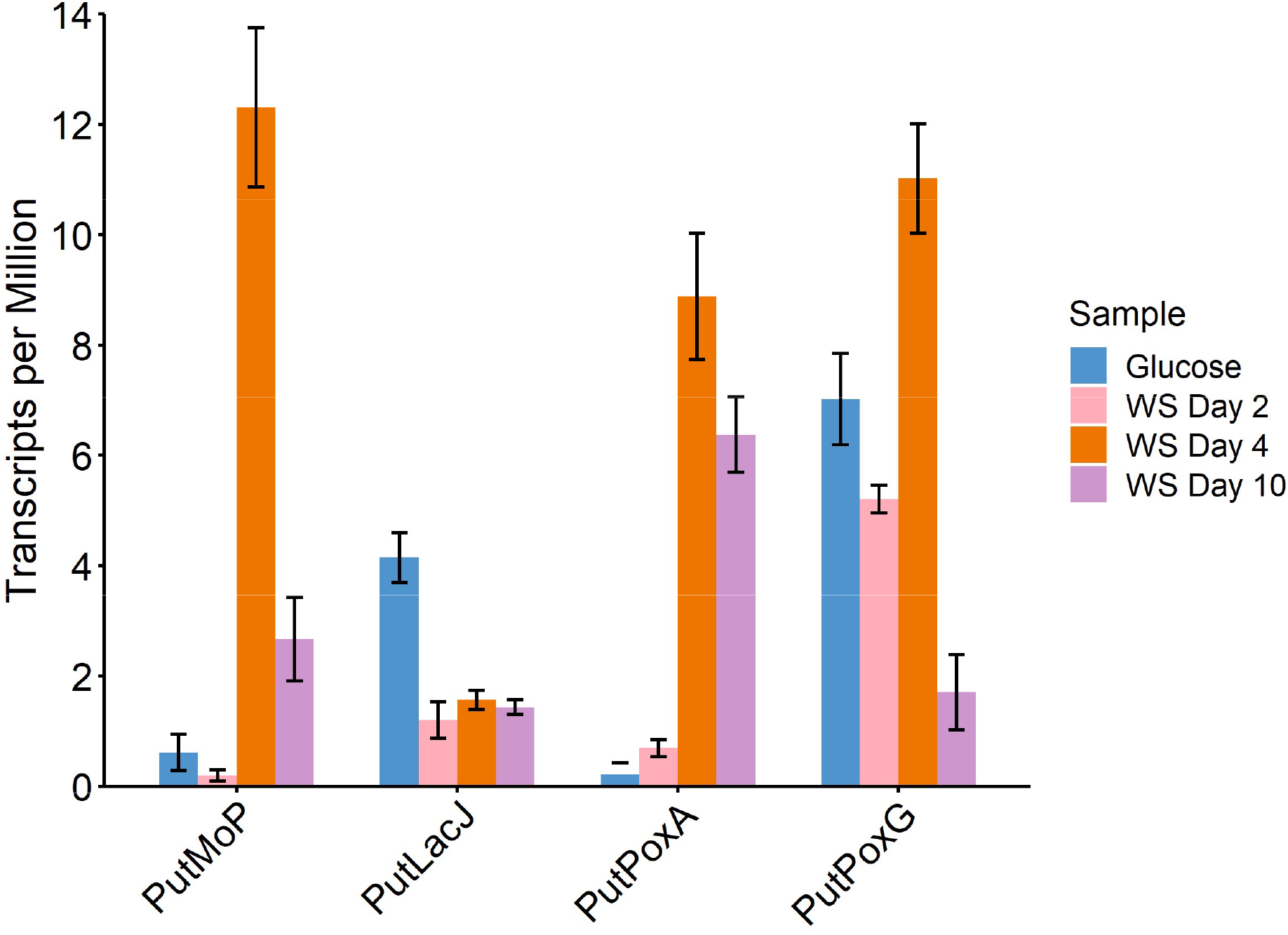
Gene Expression of Interesting Candidates. Transcripts per Million (TPM) values for each of the four candidates explored, during growth on glucose, or on day 2, 4, and 10 of growth in liquid cultures containing wheat straw (WS).

## Conclusions

*P. putredinis* NO1 was revealed here to contain a diverse repertoire of lignocellulose degrading enzymes in its genome. The newly annotated reference genome is a potentially useful resource, considering the potential of *P. putredinis* NO1 for the identification of industrially valuable enzymes (5). Among ascomycetes, *P. putredinis* NO1 exists within the 95^th^ percentile for abundant auxiliary activity gene density, implying potential specialism regarding mechanisms of lignocellulose degradation and belongs to a substantially underrepresented and underexplored lineage. Investigating CAZyme families in more detail revealed an increased capacity to target the most recalcitrant components of lignocellulose when compared to three other biomass-degrading ascomycetes. For crystalline cellulose degradation, expansions were observed in families of LPMOs and in families associated with LPMO systems. Multiple domains encoding lignin-degrading laccase proteins were also identified. Considering the context in which *P. putredinis* NO1 was identified, thriving at the late stages of a mixed microbial community grown on wheat straw, it is feasible that the genome of this fungus contains new ligninolytic activities. By utilising a strategy of searching genomic data for new enzymes with simultaneous sequence-, domain-, and structural-based approaches, multiple interesting sequences were identified.

## Experimental Methods

### Strain Isolation

*P. putredinis* NO1 was isolated from a wheat straw enrichment culture and maintained as reported previously (5).

### Genomic DNA Extraction and Sequencing

For DNA extraction, *P. putredinis* NO1 was grown in optimised media containing 10% (w/v) sucrose at 30 °C with shaking at 140 rpm for 14 days. Wet fungal biomass was washed in deionised water before pelleting in 50 mL falcon tubes at 4500 rpm for 15 minutes, and ten technical replicates of 100 mg of biomass were then prepared in 1.5 mL tubes. Fungal biomass was then digested by adding 100 µL of 1 mg mL^-1^ Chitinase from *Streptomyces griseus* (Merck) and 200 µL of 50 mM EDTA and incubating at 37 °C for 3 hours. DNA extraction was then performed with the Wizard^®^ Genomic DNA Purification Kit (Promega). Digested samples were centrifuged at 18,000 x g at 4 °C for 2 minutes and the supernatant discarded. Pellets were resuspended with 300 µL of Nuclei Lysis solution and 100 µL of Protein Precipitation solution and rotated for 5 minutes before a 5-minute incubation on ice. Samples were then centrifuged at 18,000 x g at 4 °C for 3 minutes and the supernatant transferred to fresh tubes containing 300 µL of cold isopropanol, gently mixed by inversion, and centrifuged again. The supernatant was discarded, and the pellet was washed in 70% ice cold ethanol before centrifugation followed by air drying the DNA pellet. The pellet was then resuspended in 50 µL of DNA rehydration solution with the addition of 1.5 µL of RNase solution. Samples were than incubated at 37 °C for 15 minutes followed by rehydration at 4 °C overnight. Replicate DNA samples were run on 0.75% agarose TAE gel alongside GeneRuler 1 kb Plus DNA Ladder (Thermo Scientific) at 120V for 40 minutes. The gel was then visualised in the Uvitec Gel-Documentation System to confirm the presence of long strand DNA.

Genomic DNA was subject to an additional clean up step using a 0.6:1 ratio of AMPure XP beads:sample prior to long read sequencing using the Oxford Nanopore Technologies’ (ONT) MinION system. The sequencing library was prepared using ONT’s ligation sequencing kit SQK-LSK109, as per the manufacturer’s guidelines with modifications as follows: Incubation times for end repair steps were increased from 5 minutes to 30 minutes; ligation reactions were performed at room temperature for 1 hour, and elution steps were performed at 37 °C for 15 minutes. The resulting DNA libraries were sequenced on MinION R9.4.1 flow cells with a 48-hour run time. Basecalling was performed using Guppy V 3.5.2 software.

### Genome Assembly and Annotation

Oxford Nanopore Technologies reads were filtered to those of length over 5kb with SeqKit 0.11.0 (50) before being assembled with Canu 2.0 (51). The resulting genome assembly was filtered with Tapestry 1.0.0 (52) to 39Mb, 21 contigs, before being polished with Medaka 0.11.3. Previously obtained Illumina reads were used to polish the assembly. Short read Illumina sequencing libraries were prepared using the NEBNext Ultra DNA library prep kit for Illumina (New England Biolabs), and sequenced on an Illumina HiSeq 2500, with paired end 100 bp reads, by the University of Leeds Next Generation Sequencing Facility. The Illumina reads were quality-checked with FastQC 0.11.7 (53) and adapter trimmed with Cutadapt 2.10 (54) and used for three rounds of Pilon 1.23 (55) polishing of the genome assembly. A previously obtained transcriptome assembly from NO1 grown on six lignocellulosic substrates (wheat straw, empty fruit bunches from palm oil, wheat bran, sugar cane bagasse, rice straw and kraft lignin) was used for genome annotation with FunAnnotate 1.8.1 and InterproScan 5.46 (56, 57).

### Ascomycete Genome Annotation and CAZyme Prediction

All available genome assemblies (n= 2635) of ascomycota origin were retrieved from the NCBI genome assembly database. Genome assemblies with N50 values > 1000 were retained and gene prediction was performed with FunAnnotate v1.8.1 (60), BUSCO (61), and AUGUSTUS (62), generating a final dataset of 2570 genomes. Predicted genes for each genome were annotated with the CAZyme database (v.09242921) and mean gene densities were then calculated for each taxonomic level for comparative analysis. Unique taxonomy identifiers (taxid) for each genome were retrieved from the NCBI taxonomy database using the Entrez NCBI API (58). No filtering was undertaken and a phylogenetic tree was reconstructed using ETE3 to retrieve the tree topology (get_topology) without intermediate nodes at a rank limit of genus (63) (**Figure 2**). Gene densities from annotations were mapped to the corresponding genomes on the tree. Genome metadata and annotations are available in **Supplementary File 2**.

The number and proportion of CAZyme domains in the genomes of *P. putredinis* NO1 (GCA_938049765.1), *S. boydii* (GCA_002221725.1), *T. reesei* (GCA_016806875.1), and *F. oxysporum* (GCA_023628715.1) were plotted using the ‘ggplot2’ package of R studio 3.6.3 (59, 60).

### Sequence-Based Searches for LPMOs, Laccases, and Peroxidases

The sequences for an ascomycete AA9 family LPMO and for an AA1 family laccase were obtained from the CAZy database (61). An AA9 LPMO from *Aspergillus niger* (GenBank: CAK97151.1) and an AA1 Laccase from *A. niger* (GenBank: CAK37372.1) were used. For peroxidase sequences, individual sequences for three types of reported lignin degrading peroxidases were obtained from the fPoxDB database (48). A manganese peroxidase from *Aureobasidium subglaciale* (GenBank: EJD50148.1), a lignin peroxidase from *F. oxysporum* f. sp. *lycopersici* (NCBI RefSeq: XP_018248194.1), and a versatile peroxidase from *Pyronema confluens* (Locus: PCON_11254m.01) only available from the fPoxDB database were used.

These sequences were searched against the *P. putredinis* NO1 genome protein sequences through command line BLAST with an E-value cut off of 1 x 10^-5^ (62). Results were compiled for the three classes of peroxidase.

### Domain-Based Searches for LPMOs, Laccases, and Peroxidases

Due to the lack of online databases for LPMO sequences, the genome was searched for LPMO related sequences using the Pfam AA9 HMM (31).

Sequences for basidiomycete laccases and ascomycete Multicopper oxidases were downloaded from the Laccase Engineering Database 7.1.11 (42). These were aligned using Kalign 3.0 and this alignment subsequently used to generate a bespoke Hidden Markov Model (HMM) using the HMMER 3.2.1 programme (63, 64).

Sequences for Manganese peroxidases, Lignin peroxidases and Versatile peroxidases were downloaded from the fPoxDB database (48). These were aligned and used to construct a bespoke HMM model as before.

These models were used to search the *P. putredinis* NO1 genome using HMMER 3.2.1 (64) and domain hits falling within the default significance inclusion threshold of 0.01.

### Structure-Based Searches for LPMOs, Laccases, and Peroxidases

Predicted structure for >96 % (n=9611) of coding regions in the *P. putredinis* NO1 genome were modelled using AlphaFold v2.0.0 on the VIKING computer cluster (30).

The 9611 models of coding sequences were compiled into ‘tarball’ databases and compressed into ‘.tar.gz’ files on the VIKING cluster. These files were uploaded to the PDBefold online server to search against (65). Structures for the same sequences used in sequence based searching were obtained from UniProt database if available (66), or modelled using AlphaFold v.2.00 on the VIKING computing cluster. These structures were searched against the *P. putredinis* NO1 structure database using PDBefold to identify similar structures in the *P. putredinis* NO1 genome. The ‘lowest acceptable match’ parameter was adjusted depending on the activity being searched with until coding regions not identified using sequence- or domain-based searching strategies were identified.

### *In silico* Investigation of Candidate Sequences

Sequences which were identified by structural searching solely were considered potentially interesting and warranted further investigation to attempt to elucidate function. Sequences were searched against the NCBI non-redundant protein database with default search parameters and an E-value cut off of 1 x 10^-5^ to investigate proteins with similar sequence (34). Domains were predicted using the priary amino acid sequence with the InterPro tool for domain prediction with default parameters (67). CAZyme domains were predicted with the online dbCAN prediction tool with default search parameters (7).Interesting candidate structures were further investigated with PDBefold by searching the structures against the whole PDB database to identify structurally similar proteins using a ‘lowest acceptable match’ parameter of 70% (35, 65). Secretion signals were predicted using SignalP 6.0 with default parameters (36). Altogether, this annotation information was used to investigate the potential functions of interesting sequences.

### Transcriptomic Data for Interesting Sequences

A previously published transcriptomic dataset for *P. putredinis* NO1 was used to validate expression of sequences of interest identified here during growth on lignocellulosic substrates (5). Gene expression data in transcripts per million (TPM) for all sequences identified solely by structural approaches and not by sequence- or domain-based searching and therefore considered to be interesting for all three activities explored here: LPMO, laccase, and peroxidase. Gene expression data is available in **Supplementary File 1.**

## Acknowledgements

This work was funded by the Biotechnology and Biological Sciences Research Council (BBSRC), UK (Grant BB/1018492/1, BB/P027717/1, and BB/W000695/1). CS was supported by a CASE studentship from the BBSRC Doctoral Training Programme (BB/M011151/1) with Prozomix Ltd.

This project was undertaken on the Viking Cluster, which is a high-performance compute facility provided by the University of York. We are grateful for computational support from the University of York High Performance Computing service, Viking and the Research Computing team.

Special thanks to Sally James for performing the Nanopore sequencing in her kitchen in the first weeks of the COVID-19 pandemic, and to Katherine Newling for her immense help with all bioinformatic work and my endless questions.

CJRS conceptualised the investigation carried out in this paper, extracted the genomic DNA from *P. putredinis* NO1, performed CAZyme repertoire comparison analysis, structurally modelled the *P. putredinis* NO1 genome, performed sequence-, domain-, and structure-based searches of the genome, analysed the search strategy results and was the major contributor in writing the manuscript. DRL carried out annotation of ascomycete genomes and CAZyme repertoire comparison analysis and was a major contributor to the writing of the manuscript. NCO was involved in maintaining *P. putredinis* NO1 and extraction of genomic DNA. SRJ library prepped and sequenced the *P. putredinis* NO1 genomic DNA. KN assembled the *P. putredinis* NO1 genome, performed initial annotation and aided deposition of sequence data. YL assembled the transcriptome that was used for annotation of the *P. putredinis* NO1 genome. NGSM was a contributor to the writing of the paper. SB carried out the Illumina sequencing which was used to polish the *P. putredinis* NO1 genome assembly. NCB was a major contributor to the conceptualisation and supervision of the study in addition to making a major contribution to the writing of the manuscript.

## Data Availability

The sequence data generated and analysed during the current study are available in the European Nucleotide Archive, project code PRJEB60285, secondary accession ERP145344 (https://www.ebi.ac.uk/ena/browser/view/PRJEB60285). The WGS Sequence Set for the genome assembly is available in the European Nucleotide Archive, Accession CASHTG010000000.1 (https://www.ebi.ac.uk/ena/browser/view/CASHTG010000000). The assembly is also available through the NCBI database, Accession GCA_949357655.1 (https://www.ncbi.nlm.nih.gov/datasets/genome/GCA_949357655.1/).

