## Supplemental Figures and Tables for "Whole genome structural predictions reveal hidden diversity in putative oxidative enzymes of the lignocellulose degrading ascomycete *Parascedosporium putredinis* NO1"

**<u>Supplementary Material</u>**

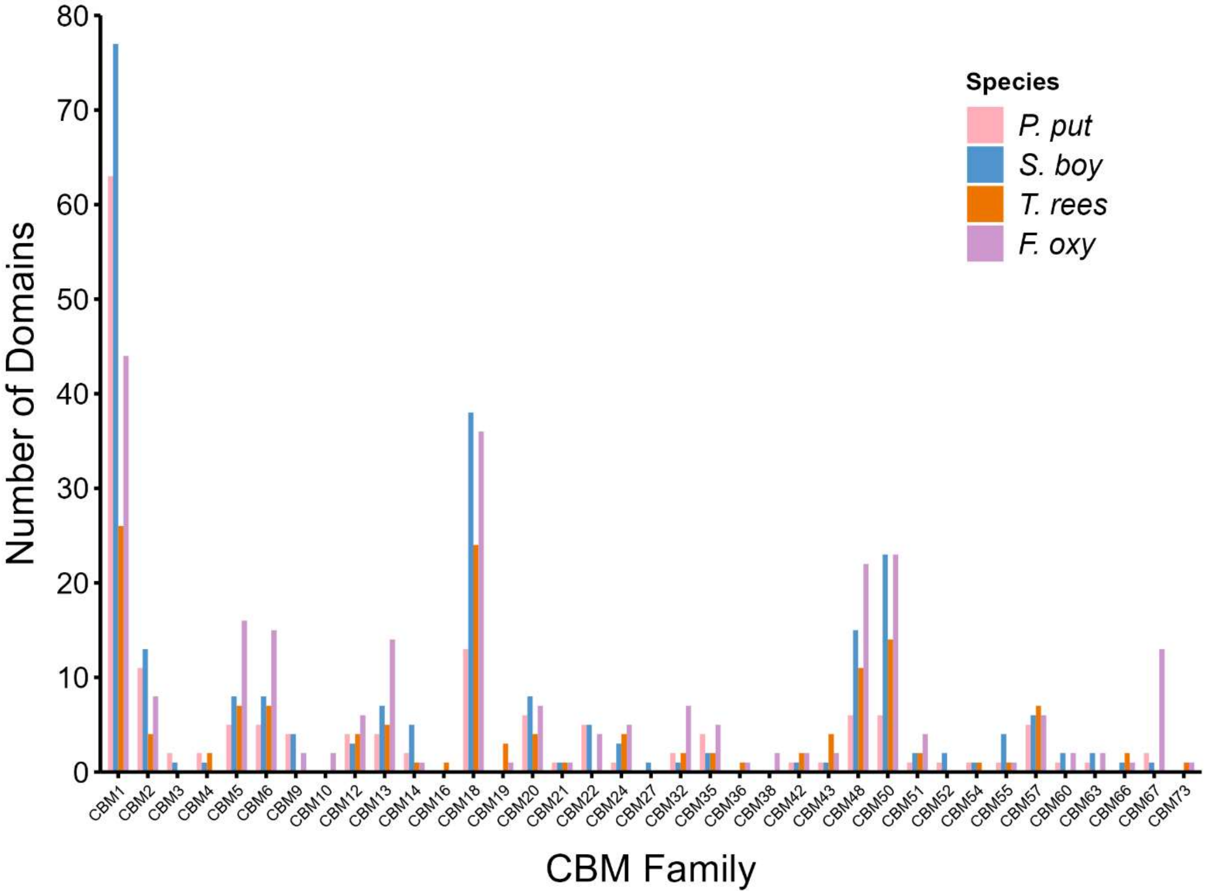


**Supplementary Figure 1. Comparison of CBM class CAZyme repertoire**. The number of CBM Class CAZyme domains of each family for four lignocellulose degrading ascomycetes; *P. putredinis* NO1, *S. boydii*, *T. reesei*, and *F. oxysporum*.

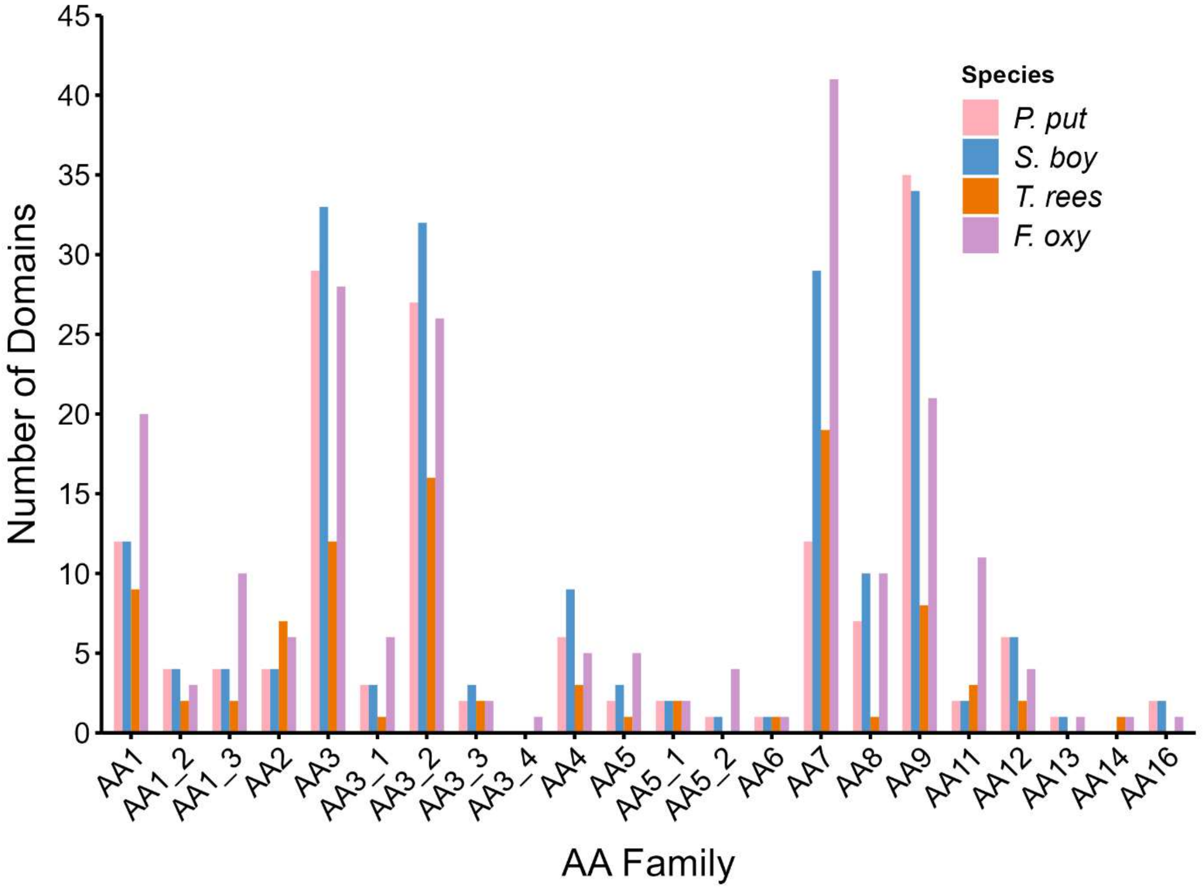


**Supplementary Figure 2. Comparison of AA class CAZyme repertoire**. The number of AA Class CAZyme domains of each family for four lignocellulose degrading ascomycetes; *P. putredinis* NO1, *S. boydii*, *T. reesei*, and *F. oxysporum*.

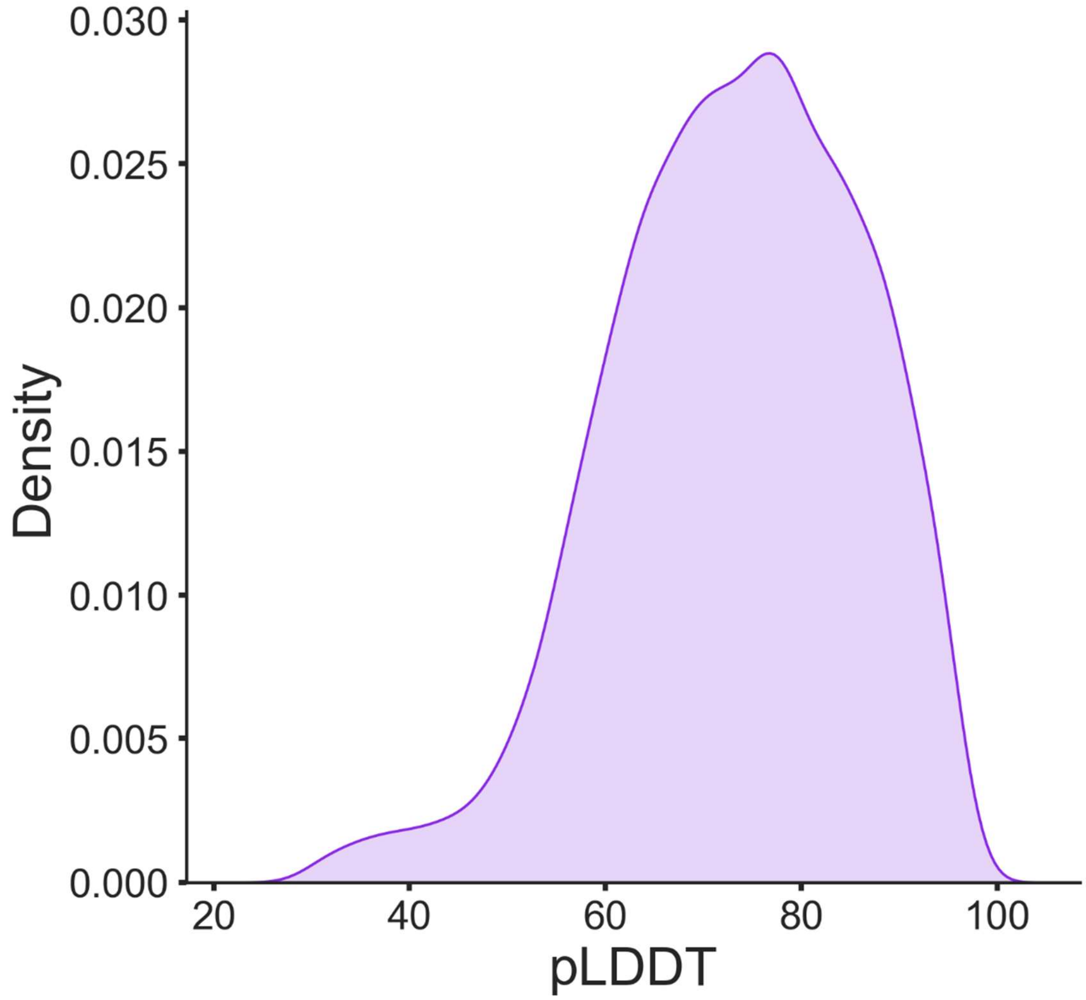


**Supplementary Figure 3. Structural Database pLDDT Score Distribution.** Density plot for structural confidence predicted local distance difference test (pLDDT) score distribution. Data shown is from 9611 structural predictions.

**Supplementary Table 1.** Coding regions of LPMO related proteins identified through genome searching approaches with the sequence of an *A. niger* AA9 LPMO (E-value cut-off = 1 x $10^{-5}$), the Pfam AA9 HMM (Significance threshold = 0.01), and the structure of the *A. niger* AA9 LPMO (Lowest percentage match = 50%).

| Coding Region | GenBank Accession | Identified by Searching Approach | | | AA9 CAZyme |
|---|---|---|---|---|---|
| | | Sequence | Domain | Structure | |
| FUN_000316-T1 | CAI7987579.1 | ✓ | ✓ | ✓ | ✓ |
| FUN_000653-T1 | CAI7987917.1 | | | ✓ | |
| FUN_000713-T1 | CAI7987978.1 | | | ✓ | |
| FUN_001321-T1 | CAI7988928.1 | ✓ | ✓ | | ✓ |
| FUN_001635-T1 | CAI7989580.1 | ✓ | ✓ | | ✓ |
| FUN_001667-T1 | CAI7989636.1 | ✓ | ✓ | ✓ | ✓ |
| FUN_001939-T1 | CAI7990224.1 | ✓ | ✓ | | ✓ |
| FUN_002327-T1 | CAI7991101.1 | ✓ | ✓ | | ✓ |
| FUN_002573-T1 | CAI7991617.1 | | ✓ | | |
| FUN_002628-T1 | CAI7991711.1 | ✓ | ✓ | ✓ | ✓ |
| FUN_002887-T1 | CAI7992274.1 | ✓ | ✓ | | ✓ |
| FUN_002890-T1 | CAI7992277.1 | | | ✓ | |
| FUN_002962-T1 | CAI7992399.1 | | | ✓ | |
| FUN_003076-T1 | CAI7992608.1 | ✓ | ✓ | ✓ | ✓ |
| FUN_003190-T1 | CAI7992922.1 | | | ✓ | |
| FUN_003419-T1 | CAI7993442.1 | | ✓ | | ✓ |
| FUN_003436-T1 | CAI7993459.1 | ✓ | ✓ | | ✓ |
| FUN_003437-T1 | CAI7993460.1 | ✓ | ✓ | ✓ | ✓ |
| FUN_003535-T1 | CAI7993628.1 | | ✓ | | |
| FUN_003783-T1 | CAI7994168.1 | | | ✓ | |
| FUN_004209-T1 | CAI7995165.1 | ✓ | ✓ | ✓ | ✓ |
| FUN_004243-T1 | CAI7995223.1 | ✓ | ✓ | ✓ | ✓ |
| FUN_004290-T1 | CAI7995316.1 | | ✓ | ✓ | ✓ |
| FUN_004866-T1 | CAI7996285.1 | ✓ | ✓ | ✓ | ✓ |
| FUN_005222-T1 | CAI7997468.1 | ✓ | ✓ | ✓ | ✓ |
| FUN_006366-T1 | CAI7999797.1 | ✓ | ✓ | ✓ | |
| FUN_006413-T1 | CAI7999893.1 | | ✓ | | |
| FUN_006553-T1 | CAI8000144.1 | | | ✓ | |
| FUN_006658-T1 | CAI8000562.1 | | ✓ | ✓ | ✓ |
| FUN_006983-T1 | CAI8001138.1 | ✓ | ✓ | ✓ | ✓ |
| FUN_007242-T1 | CAI8001774.1 | | ✓ | | |
| FUN_007243-T1 | CAI8001776.1 | | ✓ | | ✓ |
| FUN_007537-T1 | CAI8002298.1 | ✓ | ✓ | | ✓ |
| FUN_007636-T1 | CAI8002470.1 | ✓ | ✓ | ✓ | ✓ |
| FUN_007666-T1 | CAI8002525.1 | ✓ | ✓ | ✓ | |

| | | | | | |
|---|---|---|---|---|---|
| FUN_008023-T1 | CAI8003320.1 | ✓ | ✓ | ✓ | ✓ |
| FUN_008106-T1 | CAI8003467.1 | ✓ | ✓ | | |
| FUN_008107-T1 | CAI8003469.1 | | ✓ | | ✓ |
| FUN_008454-T1 | CAI8004268.1 | ✓ | ✓ | ✓ | ✓ |
| FUN_008468-T1 | CAI8004296.1 | ✓ | ✓ | | ✓ |
| FUN_008688-T1 | CAI8004812.1 | ✓ | ✓ | ✓ | ✓ |
| FUN_008818-T1 | CAI8005031.1 | ✓ | ✓ | | ✓ |
| FUN_009030-T1 | CAI7989078.1 | ✓ | ✓ | ✓ | ✓ |
| FUN_009239-T1 | CAI7992001.1 | | | ✓ | |
| FUN_009799-T1 | CAI7999895.1 | | ✓ | | ✓ |
| FUN_009919-T1 | CAI8001775.1 | ✓ | ✓ | ✓ | ✓ |
| FUN_009920-T1 | CAI8001777.1 | | ✓ | | ✓ |
| FUN_010012-T1 | CAI8003342.1 | | | ✓ | |
| FUN_010091-T1 | CAI8004298.1 | ✓ | ✓ | ✓ | ✓ |

**Supplementary Table 2.** Coding regions of laccase related proteins identified through genome searching approaches with the sequence of an *A. niger* AA1 laccase (E-value cut-off = 1 x $10^{-5}$), the bespoke laccase and multicopper oxidase HMM constructed from sequences from the laccase engineering database (Significance threshold = 0.01), and the structure of the *A. niger* AA1 laccase (Lowest percentage match = 30%).

| Coding Region | GenBank Accession | Identified by Searching Approach | | | AA1 CAZyme |
|---|---|---|---|---|---|
| | | Sequence | Domain | Structure | |
| FUN_000263-T1 | CAI7987524.1 | | ✓ | | |
| FUN_000580-T1 | CAI7987844.1 | | ✓ | | |
| FUN_000646-T1 | CAI7987911.1 | | ✓ | | |
| FUN_000758-T1 | CAI7988025.1 | ✓ | ✓ | ✓ | ✓ |
| FUN_000759-T1 | CAI7988026.1 | ✓ | ✓ | | |
| FUN_000832-T1 | CAI7988099.1 | | ✓ | | |
| FUN_001183-T1 | CAI7988671.1 | | ✓ | | |
| FUN_001583-T1 | CAI7989479.1 | | ✓ | | |
| FUN_001846-T1 | CAI7990063.1 | ✓ | ✓ | ✓ | ✓ |
| FUN_002227-T1 | CAI7990821.1 | | ✓ | ✓ | ✓ |
| FUN_002249-T1 | CAI7990863.1 | ✓ | ✓ | ✓ | |
| FUN_002296-T1 | CAI7991047.1 | ✓ | ✓ | ✓ | ✓ |
| FUN_002408-T1 | CAI7991234.1 | ✓ | ✓ | ✓ | ✓ |
| FUN_002874-T1 | CAI7992258.1 | | ✓ | | |
| FUN_003566-T1 | CAI7993680.1 | ✓ | ✓ | ✓ | ✓ |
| FUN_003732-T1 | CAI7994085.1 | | | ✓ | |
| FUN_003828-T1 | CAI7994234.1 | | ✓ | | |
| FUN_004259-T1 | CAI7995254.1 | | ✓ | | |
| FUN_004508-T1 | CAI7995689.1 | ✓ | ✓ | ✓ | ✓ |
| FUN_004577-T1 | CAI7995802.1 | ✓ | ✓ | ✓ | ✓ |
| FUN_004616-T1 | CAI7995870.1 | | ✓ | | |
| FUN_004739-T1 | CAI7996089.1 | | ✓ | | |
| FUN_005070-T1 | CAI7996980.1 | ✓ | ✓ | ✓ | ✓ |
| FUN_005132-T1 | CAI7997298.1 | | ✓ | ✓ | |
| FUN_005520-T1 | CAI7998008.1 | | ✓ | | |
| FUN_006244-T1 | CAI7999594.1 | | ✓ | | |
| FUN_006620-T1 | CAI8000270.1 | | ✓ | | |
| FUN_006720-T1 | CAI8000684.1 | | ✓ | | |
| FUN_007228-T1 | CAI8001746.1 | | ✓ | | |
| FUN_007508-T1 | CAI8002246.1 | | ✓ | | |
| FUN_008329-T1 | CAI8004041.1 | | ✓ | | |
| FUN_009491-T1 | CAI7995256.1 | | ✓ | | |

**Supplementary Table 3.** Coding regions of peroxidase related proteins identified through genome searching approaches with the sequences of an MnP from *A. subglaciale*, LiP from *F. oxysporum*, and VP from *P. confluens* (E-value cut-off = $1 \times 10^{-5}$), the bespoke peroxidase HMM contricted from MnP, LiP, and VP seqeunces in the fPoxDB database (Significance threshold = 0.01), and the structure of the same three peroxidases used for seqeunce searches (Lowest percentage match = 30%).

| Coding Region | GenBank Accession | Identified by Searching Approach | | | AA2 CAZyme |
|---|---|---|---|---|---|
| | | Sequence | Domain | Structure | |
| FUN_002995-T1 | CAI7992466.1 | | | ✓ | |
| FUN_003542-T1 | CAI7993642.1 | | | ✓ | |
| FUN_003618-T1 | CAI7993895.1 | | | ✓ | |
| FUN_004484-T1 | CAI7995643.1 | | | ✓ | |
| FUN_004903-T1 | CAI7996357.1 | | ✓ | | ✓ |
| FUN_004941-T1 | CAI7996699.1 | ✓ | ✓ | ✓ | ✓ |
| FUN_005340-T1 | CAI7997684.1 | ✓ | ✓ | ✓ | ✓ |
| FUN_008413-T1 | CAI8004205.1 | | | ✓ | |
| FUN_008923-T1 | CAI7988420.1 | | | ✓ | |
| FUN_009329-T1 | CAI7993214.1 | | | ✓ | |

**Supplementary File 1. Gene expression of interesting sequences.** Sequences identified solely by structural searching approaches were considered interesting and were searched for in transcriptomic data from triplicate cultures of *P. putredinis* NO1 grown on glucose or grown on wheat straw with samples taken at days 2, 4, and 10. Gene expression values are presented in Transcripts per million (TPM).

**Supplementary File 2. Ascomycete genome annotations.** Annotations of all ascomycete genomes used in this analysis (n = 2570).
